# *Lactobacillus rhamnosus* Lcr35^®^ as an effective treatment for preventing *Candida albicans* infection in preclinical models: first mechanistical insights

**DOI:** 10.1101/612481

**Authors:** Cyril Poupet, Taous Saraoui, Philippe Veisseire, Muriel Bonnet, Caroline Dausset, Marylise Gachinat, Olivier Camarès, Christophe Chassard, Adrien Nivoliez, Stéphanie Bornes

**Affiliations:** Université Clermont Auvergne, INRA, VetAgro Sup, UMRF,15000, Aurillac, France; biose Industrie, 24 avenue Georges Pompidou, Aurillac, France

**Keywords:** *Lactobacillus rhamnosus* Lcr35^®^, *Candida albicans*, *Caenorhabditis elegans*, prophylaxis, immune response

## Abstract

The increased recurrence of *Candida albicans* infections is associated with greater resistance to antifungal drugs. This involves the establishment of alternative therapeutic protocols such as the probiotic microorganisms whose antifungal potential has already been demonstrated using preclinical models (cell cultures, laboratory animals). Understanding the mechanisms of action of probiotic microorganisms has become a strategic need for the development of new therapeutics for humans. In this study, we investigated the prophylactic anti-*Candida albicans* properties of *Lactobacillus rhamnosus* Lcr35^®^ using the *in vitro* Caco-2 cells model and the *in vivo Caenorhabditis elegans* model. On Caco-2 cells, we showed that the strain Lcr35^®^ significantly inhibited the growth of the pathogen (~2 log CFU.mL^−1^) and its adhesion (150 to 6,300 times less). Moreover, on the top of having a prolongevity activity in the nematode, Lcr35^®^ protects the animal from the fungal infection even if the yeast is still detectable in its intestine. At the mechanistic level, we noticed the repression of genes of the p38 MAPK signaling pathway and genes involved in the antifungal response induced by Lcr35^®^ suggesting that the pathogen no longer appears to be detected by the worm immune system. However, the DAF-16 / FOXO transcription factor, implicated in the longevity and antipathogenic response of *C. elegans*, is activated by Lcr35^®^. These results suggest that the probiotic strain acts by stimulating its host via DAF-16, but also by suppressing the virulence of the pathogen.

## 1 Introduction

*Candida albicans* is a commensal yeast found in the gastrointestinal and urogenital tracts (1,2), responsible for various infections ranging from superficial infections affecting the skin to life-threatening systemic infections i.e. candidemia (3). Its pathogenicity is based on several factors as the formation of biofilms, thigmotropism, adhesion and invasion of host cells, secretion of hydrolytic enzymes (3) and a transition from yeast to hyphal filaments facilitating its spread (4,5).

There is an increase in the number of fungal infections mainly due to the increase in resistance to drugs (6,7) and to the limited number of available antifungals, some of which are toxic (8). In addition, it is very common that antifungal treatment destabilizes more or less severely the host commensal microbiota, leading to dysbiosis (9). This state creates a favorable situation for the establishment of another pathogen or a recurrence. Besides, because of the presence of similarities between yeasts and human cells (i.e. eukaryotic cells), the development of novel molecules combining antifungal activity and host safety was particularly complicated (8). These different elements demonstrate the need to develop new therapeutic strategies aimed at effectively treating a fungal infection while limiting the health risks for the host in particular by preserving the integrity of its microbiota. The use of probiotics in order to cure candidiasis or fungal-infection-related dysbiosis is part of theses novel strategies (10–12). The World Health Organization (WHO) and the Food and Agriculture Organization of the United Nations (FAO) defines probiotics as “live microorganisms, which, when administered in adequate amounts, confer a health benefit on the host” (13). Under this appellation of probiotic, a wide variety of microbial species is found within both prokaryotes and eukaryotes (yeasts like *Saccharomyces*) although these are mainly lactic bacteria such as the genera *Lactobacillus* and *Bifidobacterium* (14). Nowadays, a new name is increasingly used to replace the term probiotic: live biotherapeutic products (LBP). These LBP are biological products containing live biotherapeutic microorganisms (LBM) used to prevent, treat or cure a disease or condition of human beings, excluding vaccines (15).

In this issue, we focus on *Lactobacillus rhamnosus* Lcr35^®^, a Gram-positive bacterium commercialized by biose^®^ as a pharmaceutical product for more than 60 years for preventive and curative gastrointestinal and gynecological indications. Lcr35^®^ is a well-known probiotic strain whose *in vitro* and *in vivo* characteristics are widely documented (16–23). Nivoliez *et al.* demonstrated the probiotic properties of the native strain such as resistance to gastric acidity and bile stress, lactic acid production. Under its commercial formulations, Lcr35^®^ strain has the ability to adhere on intestinal (Caco-2, HT29-MTX) and vaginal (CRL -2616) epithelial cells. The inhibition of the pathogens’ adhesion to the intestinal cells by Lcr35^®^ has not been investigated by the authors. This study has also shown that Lcr35^®^ leads to a strong inhibition of vaginal (*Candida albicans*, *Gardnerella vaginalis*) and intestinal (enterotoxigenic and enteropathogenic *Escherichia coli* (ETEC, EPEC), *Shigella flexneri*) pathogens (24). Although these probiotic and antimicrobial effects have been observed during clinical trials but we know little about the molecular mechanisms underlying these properties. Randomized trials conducted in infants and children have shown that preventive intake of probiotics has a positive impact on the development of infectious or inflammatory bowel diseases by maintaining the balance of the microbiota (Isolauri *et al.* 2002). *In vitro* as well as *in vivo* studies, using preventive approaches, have revealed certain mechanisms of action of probiotics (26).

Up to now, most probiotics used in both food and health applications are selected and characterized on the basis of their *in vitro* properties (27) before being tested on complex *in vivo* models (murine models) and in human clinical trials. The *in vitro* are used mainly for ethical and cost issues (28) but also allow experimentations under defined and controlled conditions. As a result, some strains meeting the criteria for *in vitro* selection no longer respond *in vivo* and vice versa (29). This fact reinforces the idea that *in vitro* and *in vivo* tests are complementary and necessary for the most reliable characterization of probiotic properties.

Here we propose to use both the *in vitro* Caco-2 cells culture and the invertebrate host *Caenorhabditis elegans* as an *in vivo* model to investigate the microorganism – microorganism – host interactions. Caco-2 cells are a well characterized enterocyte-like cell line. They are a reliable *in vitro* system to study the adhesion capacity of lactobacilli as well as their probiotic effects, such as protection against intestinal injury induced by pathogens (30,31). Nevertheless, the use of *in vivo* models, allowing to get closer to the complex environment of the human body, is inevitable in the case of a mechanistic study. Indeed, while rudimentary models such as *Caenorhabditis elegans*, or *Drosophila* exhibit obvious benefits for (large) screening purposes, they are also not devoid of relevance in deciphering more universal signaling pathways, even related to mammalian innate immunity (32). With its many genetic and protein homologies with human beings (33), *C. elegans* has become the ideal laboratory tool for physiological as well as mechanistic studies. The roundworm has already been used to study the pathogenicity mechanisms of *Candida albicans.* Pukkila-Worley *et al*. have demonstrated a rapid antifungal response with the overexpression of antimicrobials encoding genes such as *abf-2*, *fipr-22, fipr-23*, *cnc-7*, *thn-1* and chitinases (*cht-1* and T19H5.1) or detoxification enzymes (*oac-31*, *trx-3*). It has also been shown that *C. albicans* hyphal formation is a key virulence factor who modifies the gene expression in the *C. elegans* killing assay (34). Some of these genes are notably dependent on the highly conserved p38 MAPK signaling pathway (35). Several recent studies have established that the transition from yeast morphology to hyphal form was largely dependent on environmental parameters. It is also controlled by genetic factors such as eIF2 kinase Gcn2 (36) or SPT20 (37) whose mutations induce a decrease in virulence of the pathogen and an enhanced survival of the host. However, few studies have been conducted with the nematode on the use of probiotic microorganisms for the treatment of *C. albicans* fungal infection (38).

In this context, the aim of this study was to evaluate the effect of *Lactobacillus rhamnosus* Lcr35^®^ strain to prevent a fungal infection due to *C. albicans* using the *in vitro* cellular model Caco-2 and the *in vivo* model *C. elegans*. In order to overcome the experimental limits of the *in vitro* model, we conducted the mechanistic study solely on the *C. elegans* model. The worm survival and gene expression, in response to the pathogen and/or the probiotic, were evaluated.

## 2 Material and methods

### 2.1 Microbial strains and growth conditions

*Escherichia coli* OP50 strain was provided by the *Caenorhabditis* Genetics Center (Minneapolis, MN, USA) and was grown on Luria Broth (LB, MILLER’S Modification) (Conda, Madrid, Spain) at 37 °C overnight. *Lactobacillus rhamnosus* Lcr35^®^ strain was provided by biose^®^ (Aurillac, France) and was grown in de Man, Rogosa, Sharpe (MRS) broth (bioMérieux, Marcy l’Etoile, France) at 37 °C overnight. *Candida albicans* ATCC 10231 was grown in Yeast Peptone Glucose (YPG) broth pH 6.5 (per L: 10 g yeast extract, 10 g peptone, 20 g glucose) at 37 °C for 48 hours. Microbial suspensions were spin down for 2 minutes at 1,500 rpm (Rotofix 32A, Hettich Zentrifugen, Tuttlingen, Germany) and washed with M9 buffer (per L: 3 g KH_2_PO_4_, 6 g Na_2_HPO_4_, 5 g NaCl, 1 mL 1 M MgSO_4_) in order to have a final concentration of 100 mg.mL^−1^.

### 2.2 Influence of Lcr35^®^ on *Candida albicans* growth and on *Candida albicans* biofilm formation on Caco-2 cells monolayer

Growth inhibition of *C. albicans* by the probiotic strain Lcr35^®^ was examined using the human colorectal adenocarcinoma cell line Caco-2 (39). Caco-2 cells were grown in Dulbecco modified Eagle’s minimal essential medium (DMEM, LIFE TECHNOLOGIE, Villebon-sur-Yvette, France) supplemented with 20% inactivated fetal calf serum (LIFE TECHNOLOGIE, Villebon-sur-Yvette, France) at 37 °C with a 5% CO_2_ in air atmosphere. For the assays, the cells were seeded at a concentration of 3.5×10^5^ cells.well^−1^ in 24-well plates (DUTSCHER, Brumath, France) and placed in growth conditions for 24 hours. Microbial strains were grown according to Nivoliez *et al.* (24). After growth, cell culture medium is removed and replaced by 1 mL of DMEM and 250 μL of Lcr35^®^ culture (10^8^ CFU.mL^−1^) in each well and incubated for 24 hours. 250 μL of *C. albicans* culture at different concentrations (10^7^, 10^6^, 10^5^, 10^4^, 10^3^ and 10^3^ CFU.mL^−1^) are added in each well. After incubation for 24 and 48 hours, the inhibition of *C. albicans* by Lcr35^®^ is evaluated. 100 μL of suspension is taken from each of the wells and the number of viable bacteria and/or yeasts were determined by plating serial dilutions of the suspensions onto MRS or Sabouraud agar plates. For the measurement of *C. albicans* biofilm formation, after incubation for 48 hours, the wells were washed twice with 0.5 mL of PBS and cells harvested with 1 mL of trypsin at 37 °C. As for the inhibition assay, the number of viable bacteria or/and yeasts were determined by plating serial dilutions of the suspensions onto MRS or Sabouraud agar plates. The plates are incubated at 37 °C for 72 hours (MRS) or 48 hours (Sabouraud). Each assay, performed three times independently, contains two technical replicates.

### 2.3 *Caenorhabditis elegans* maintenance

*Caenorhabditis elegans* N2 (wild-type) and TJ356 (*daf-16p::daf-16a/b::GFP* + *rol-6(su1006)*) strains were acquired from the *Caenorhabditis* Genetics Center (Minneapolis, MN). The nematodes were grown and maintained at 20 °C on Nematode Growth Medium (NGM) (per L: 3 g NaCl; 2.5 g peptone; 17 g agar; 5 mg cholesterol; 1 mM CaCl_2_; 1 mM MgSO_4_, 25 mL 1 M potassium phosphate buffer at pH 6) plates, supplemented with yeast extract (4 g.L^−1^) (NGMY) and seeded with *E. coli* OP50 (40).

### 2.4 *Caenorhabditis elegans* synchronization

In order to avoid variation in results due to age differences, a worm synchronous population is required. Gravid worms were washed off using M9 buffer and spin down for 2 minutes at 1,500 rpm. 5 mL of worm bleach (2.5 mL of M9 buffer, 1.5 mL of bleach, 1 mL of sodium hydroxide 5M) was added to the pellet and vigorously shaken until adult worm body disruption. The action of worm bleach was stopped by adding 20 mL of M9 buffer. Eggs suspension was then spun down for 2 minutes at 1,500 rpm and washed twice with 20 mL of M9 buffer. Eggs were allowed to hatch under slow agitation at 25 °C for 24 hours in about 20 mL of M9 buffer. L1 larvae were then transferred on NGMY plates seeded with *E. coli* OP50 until they reach L4 / young adult stage.

### 2.5 Body size

Individual adult worms were photographed using an Evos FL microscope (Invitrogen, 10X magnification). After reaching L4 stage, they were transferred on NGMY plates previously seeded with the probiotic strain Lcr35^®^ and their size were measured daily for three days. Length of worm body was determined by using ImageJ software as described by Mörck and Pilon (2006) (41) and compared to OP50-fed worms. At least 10 nematodes per experiment were imaged on at least three independent experiments.

### 2.6 *Caenorhabditis elegans* lifespan assay

Synchronous L4 worms were transferred on NGMY with 0.12 mM 5-fluorodeoxyuridine FUdR (Sigma, Saint-Louis, USA) and seeded with 100 μL of the 100 mg.mL^−1^ microbial strain (~50 worms per plate). The plates were kept at 20 °C and live worms were scored each day until the death of all animals. An animal was scored as dead when it did not respond to a gentle mechanical stimulation. This assay was performed as three independent experiments with three plates per condition.

### 2.7 Effects of *Lactobacillus rhamnosus* Lcr35^®^ on candidiasis in *Caenorhabditis elegans*

Sequential feeding with Lcr35^®^ and *C. albicans* were induced in *C. elegans* in all experiments (preventive assays). As control groups, a monotypic contamination was induced in *C. elegans* by inoculation only of *C. albicans*, Lcr35^®^ or *E. coli* OP50.

#### 2.7.1 Preparation of plates containing probiotic bacteria or pathogen yeasts

100 μL of Lcr35^®^ or *E. coli* OP50 suspension (100 mg.mL^−1^) was spread on NGMY + 0.12 mM FUdR plates and incubated at 37 °C overnight. Concerning *C. albicans* strains, 100 μL of suspension were spread on Brain Heart Infusion BHI (Biokar diagnostics, Beauvais, France) + 0.12 mM FUdR plates and incubated at 37 °C overnight.

#### 2.7.2 Survival assay: preventive treatment

The survival assay was performed according to de Barros *et al.* 2018 (38), with some modifications. During a preventive treatment, young adult worms were placed on plates containing Lcr35^®^, at 20 °C for different times (2, 4, 6 and 24 hours). Next, worms are washed with M9 buffer to remove bacteria prior being placed on *C. albicans* plates for 2 hours at 20 °C. Infected nematodes were washed off plates using M9 buffer prior to be transferred into a 6-well microtiter plate (about 50 worms per well) containing 2 mL of BHI / M9 (20% / 80%) + 0.12 mM FUdR liquid assay medium per well and incubated at 20 °C. For the control groups (i.e. *E. coli* OP50 + *C. albicans*, *E. coli* OP50 only, Lcr35^®^ only and *C. albicans* only), worms were treated in the same way. Nematodes were observed daily and were considered dead when they did not respond to a gentle mechanical stimulation. This assay was performed as three independent experiments containing three wells per condition.

### 2.8 Colonization of *C. elegans* intestine by *C. albicans*

In order to study worm’s gut colonization by the pathogen *C. albicans*, a fluorescent staining of the yeast was performed. The yeast was stained with rhodamine 123 (Yeast Mitochondrial Stain Sampler Kit, Invitrogen, Eugene, USA) according to the manufacturer’s instructions. A fresh culture of *C. albicans* was done in YPG broth as described before, 1.6 μL of rhodamine 123 at 25 mM is added to 1 mL of *C. albicans* suspension and incubated at room temperature in the dark for 15 minutes. The unbound dye is removed by centrifugation (14,000 rpm for 5 minutes at 4 °C) (Beckman J2-MC Centrifuge, Beckman Coulter, Brea, USA) and washed with 1 mL of M9 buffer. Subsequently, the nematodes are fed on *E. coli* OP50 or Lcr35^®^ on NGMY plates for 4 hours and then with labeled *C. albicans* on BHI plates for 72 hours. The nematodes are then visualized using a 100X magnification fluorescence microscope (Evos FL, Invitrogen).

### 2.9 RNA isolation and RT-quantitative PCR

About 10,000 worms were harvested from NGMY plates with M9 buffer. Total RNA was extracted by adding 500 μL of TRIzol reagent (Ambion by life technologies, Carlsbad, USA). Worms were disrupted by using a Precellys (Bertin instruments, Montigny-le-Bretonneux, France) and glass beads (PowerBead Tubes Glass 0.1mm, Mo Bio Laboratories, USA). Beads were removed by centrifugation at 14,000 rpm for 1 minute (Eppendorf^®^ 5415D, Hamburg, Germany), and 100 μL of chloroform were added to the supernatant. Tubes were vortexed for 30 seconds and incubated at room temperature for 3 minutes. The phenolic phase was removed by centrifugation at 12,000 rpm for 15 minutes at 4 °C. The aqueous phase was treated with chloroform as previously. RNA was precipitated by adding 250 μL of isopropanol for 4 minutes at room temperature and spin down at 12,000 rpm for 10 minutes (4 °C). The supernatant was discarded and the pellet was washed with 1,000 μL of 70% ethanol. The supernatant was discarded after centrifugation at 14,000 rpm for 5 minutes (4 °C) and the pellet was dissolved into 20 μL of RNase-free water. RNA was reverse-transcribed using High-Capacity cDNA Archive kit (Applied Biosystems, Foster City, USA), according to the manufacturer’s instructions. For real-time qPCR assay, each tube contained 2.5 μL of cDNA, 6.25 μL of Rotor-Gene SYBR Green Mix (Qiagen GmbH, Hilden, Germany), 1.25 μL of 10 μM primers (reported in Table 1) (Eurogentec, Seraing, Belgium) and 1.25 μL of water. All samples were run in triplicate. Rotor-Gene Q Series Software (Qiagen GmbH, Hilden, Germany) was used for the analysis. In our study, two reference genes, *cdc-42* and Y45F10D.4, were used in all the experimental groups. The Quantification of gene-of-interest expression (E_GOI_) was performed according to Hellemans *et al.* formula (42):

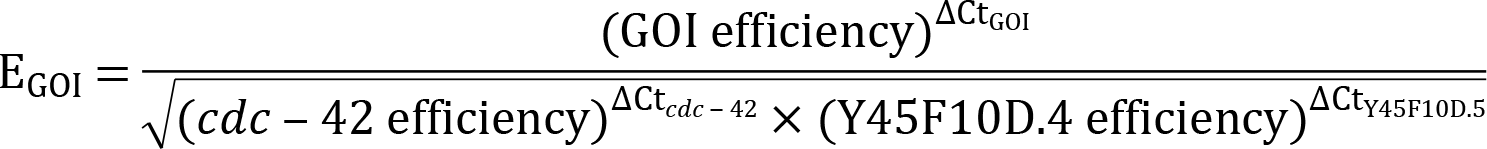

**Table 1:**
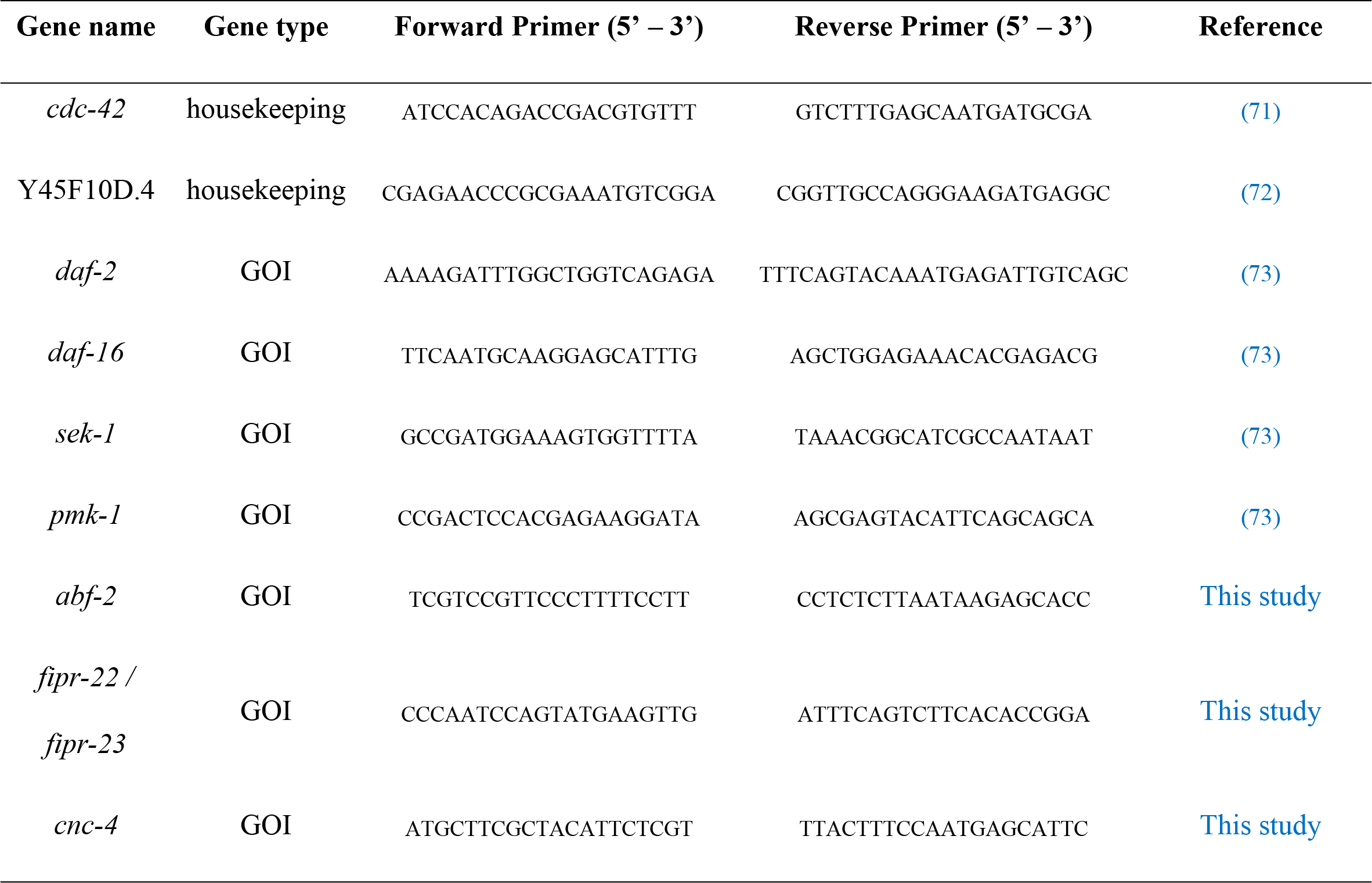
Targeted *C. elegans* genes primers for qPCR analysis. GOI: Gene of Interest

### 2.10 Statistical analysis

Data are expressed as the mean ± standard deviation.

*C. elegans* survival assay was examined by using the Kaplan-Meier method, and differences were determined by using the log-rank test with R software version 3.5.0 (43), *survival* (44) and *survminer* (45) packages. For *C. albicans* growth inhibition and biofilm formation, *C. elegans* growth and gene expression of the genes analyzed, differences between conditions were determined by a two-way ANOVA followed by a Fisher’s Least Significant Difference (LSD) post hoc test using GraphPad Prism version 7.0a for Mac OS X (GraphPad Software, La Jolla, California, USA). A *p*-value ≤ 0.05 was considered as significant.

### 2.11 DAF-16 nuclear localization

DAF-16 nuclear localization was followed as described by Fatima *et al.* 2014 (46) using transgenic TJ-356 worms (DAF-16::GFP). Once adults, worms are exposed to single strain: *E. coli* OP50, Lcr35^®^ or *C. albicans* for 2, 4, 6, 24 and 76 hours at 20 °C. A preventive approach was also conducted: worms were put in the presence of *E. coli* OP50 or Lcr35^®^ for 4 hours then *C. albicans* for 2 hours. The nematodes were subsequently photographed 2, 4, 6 and 24 hours after infection. The translocation of DAF-16::GFP was scored by assaying the presence of GFP accumulation in the *C. elegans* cell nuclei, using a 40X magnification fluorescence microscope (Evos FL, Invitrogen).

## 3 Results

### 3.1 Anti-*Candida albicans* effects of Lcr35^®^ on Caco-2 cell monolayer

#### 3.1.1 Growth inhibition of the yeast

In the presence of Caco-2 cells, regardless of the concentration of the inoculum (from 10^2^ to 10^7^ CFU.mL^−1^), *C. albicans* grew to concentrations that ranged from 7.48 ± 0.39 to 7.83 ± 0.34 log CFU.mL^−1^ after 48 hours of incubation. Similar *C. albicans* growth was measured in the absence of human cells (data not shown). When prophylactic treatment was used, i.e. when the Caco-2 cells were pre-incubated with the probiotic Lcr35^®^, we observed an antifungal activity against *C. albicans*. Indeed, the bacterium induced a significant inhibition of the yeast of 2 log CFU.mL^−1^ which then reached a concentration ranging from 5.40 ± 0.07 to 6.05 ± 0.25 log CFU.mL^−1^. Two different inhibition profiles were observed after 48 h. On one hand, when the inoculum was highly concentrated (7 log CFU.mL^−1^), we observed a decrease in the yeast population which is a sign of cell death. On the other hand, when the inoculum was less concentrated (2 to 4 log CFU.mL^−1^), we noticed that the yeast was able to grow although its growth seemed to stop between 5.32 ± 0.36 and 5.51 ± 0.14 log CFU.mL^−1^ (Table 2).

**Table 2:** Monitoring of *Candida albicans* growth in the presence of Lcr35^®^ on Caco-2 cells monolayer. Results are expressed as log_10_ CFU.mL^−1^ of yeasts alone (controls) in co-incubation with Lcr35^®^ (mean ± standard deviation). Comparison between conditions with and without Lcr35^®^ was performed using a two-way ANOVA followed by a Fisher’s LSD post hoc test (p < 0.05: * ; p < 0.01: ** ; p < 0.001 : *** ; p < 0.0001 : ****)

|  |  | Length of incubation (hours) |  |  |
| --- | --- | --- | --- | --- |
| Concentration of *Candida albicans* inocula (CFU.mL<sup>-1</sup>) | With or without Lcr35® | 0 | 24 | 48 |
| 10<sup>7</sup> | with | 7.25 ± 0.51 | 6.39 ± 0.73 | 6.05 ± 0.25 **** |
|  | without | 6.77 ± 0.10 | 7.29 ± 0.23 | 7.78 ± 0.41 |
| 10<sup>6</sup> | with | 5.85 ± 0.25 | 5.47 ± 0.12 * | 5.73 ± 0.09 *** |
|  | without | 5.76 ± 0.18 | 7.42 ± 0.27 | 7.69 ± 0.20 |
| 10<sup>5</sup> | with | 4.77 ± 0.41 | 5.01 ± 0.12 ** | 5.49 ± 0.04 **** |
|  | without | 4.60 ± 0.28 | 7.60 ± 0.69 | 7.83 ± 0.34 |
| 10<sup>4</sup> | with | 3.69 ± 0.21 | 4.92 ± 0.54 | 5.51 ± 0.14 * |
|  | without | 3.72 ± 0.13 | 7.09 ± 0.59 | 7.48 ± 0.39 |
| 10<sup>3</sup> | with | 2.56 ± 0.34 | 3.59 ± 0.25 | 5.51 ± 0.16 **** |
|  | without | 2.30 ± 0.17 | 6.60 ± 0.28 | 7.93 ± 0.45 |
| 10<sup>2</sup> | with | 1.34 ± 0.31 | 3.18 ± 0.76 | 5.32 ± 0.36 *** |
|  | without | 1.34 ± 0.38 | 6.18 ± 1.01 | 7.80 ± 0.27 |

#### 3.1.2 Inhibition of the yeast’s biofilm formation

The ability of a pathogen to form a biofilm is an important step in facilitating its systemic dissemination in the host tissue. After 48 hours of incubation, the *C. albicans* biofilm contained between 5.78 log CFU.mL^−1^ (inoculum at 10^2^ CFU.mL^−1^) and 8.69 log CFU.mL^−1^ of yeasts (inoculum at 10^7^ CFU.mL^−1^). However, since the cells were pre-exposed to Lcr35^®^ and for the same *C. albicans* inocula, we observed a significant decrease in the amount of yeasts in the biofilm: 4.32 to 5.16 log CFU.mL^−1^, which corresponded to an inhibition ranging from 1.46 to 3.53 log. The strongest inhibition was observed in the case where the inoculum of *C. albicans* was the most concentrated (Fig 1).

**Fig 1:**
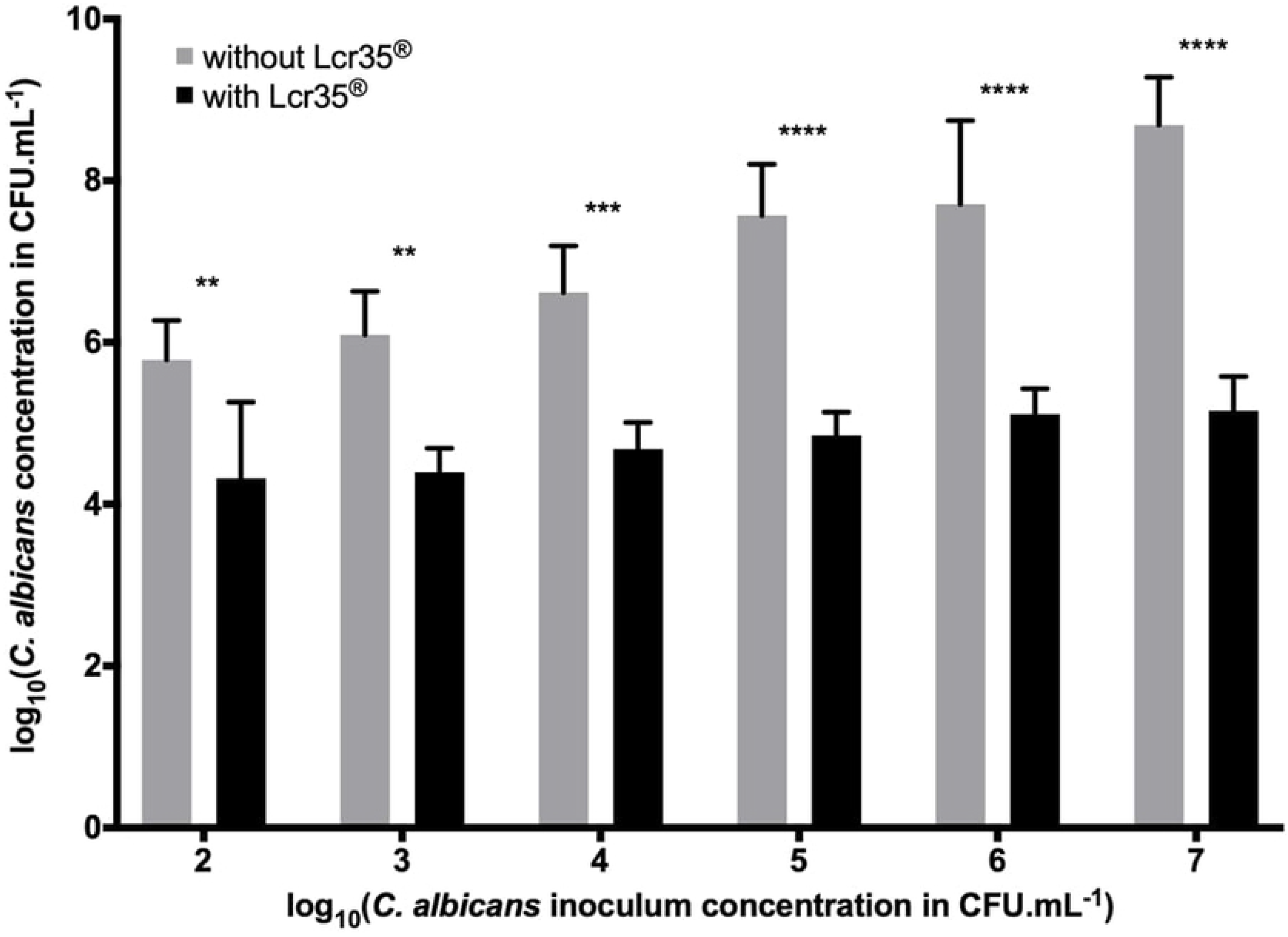
Determination of the *C. albicans* biofilm formation in presence of Lcr35^®^ (10^8^ CFU.mL^−1^) or not onto Caco-2 cells monolayer (mean ± standard deviation). Different concentrations of yeasts were tested then the amount present in the biofilm was evaluated after 48 hours of incubation. Comparison between conditions with and without Lcr35^®^ was performed using a two way ANOVA followed by a Fisher’s LSD post hoc (p < 0.05: * ; p < 0.01: ** ; p < 0.001 : *** ; p < 0.0001 : ****)

### 3.2 Effects of Lcr35^®^ on *C. elegans* physiology

#### 3.2.1 Lcr35^®^ extends *C. elegans* lifespan

We investigated the effects on *C. elegans* lifespan induced by either the pathogenic yeast *C. albicans* or the probiotic Lcr35^®^. Feeding adult nematodes with the probiotic strain resulted in a significant increase of the mean lifespan compared to OP50-fed worms (p = 3.56 .10^−6^) evolving from 7 to 10 days (+ 42.9%) whereas *C. albicans* had no impact on *C. elegans* mean lifespan. On the other hand, when *C. albicans* was used as a feeding source, worms displayed a significant reduced lifespan (p = 1.27 .10^−5^) which dropped from 16 to 14 days (−12.5%). Lcr35^®^ did not increase the worm’s longevity compared to OP50 (Fig 2). These results showed that the probiotic strain ameliorated the mean lifespan without increasing the life expectancy of the worm.

**Fig 2:**
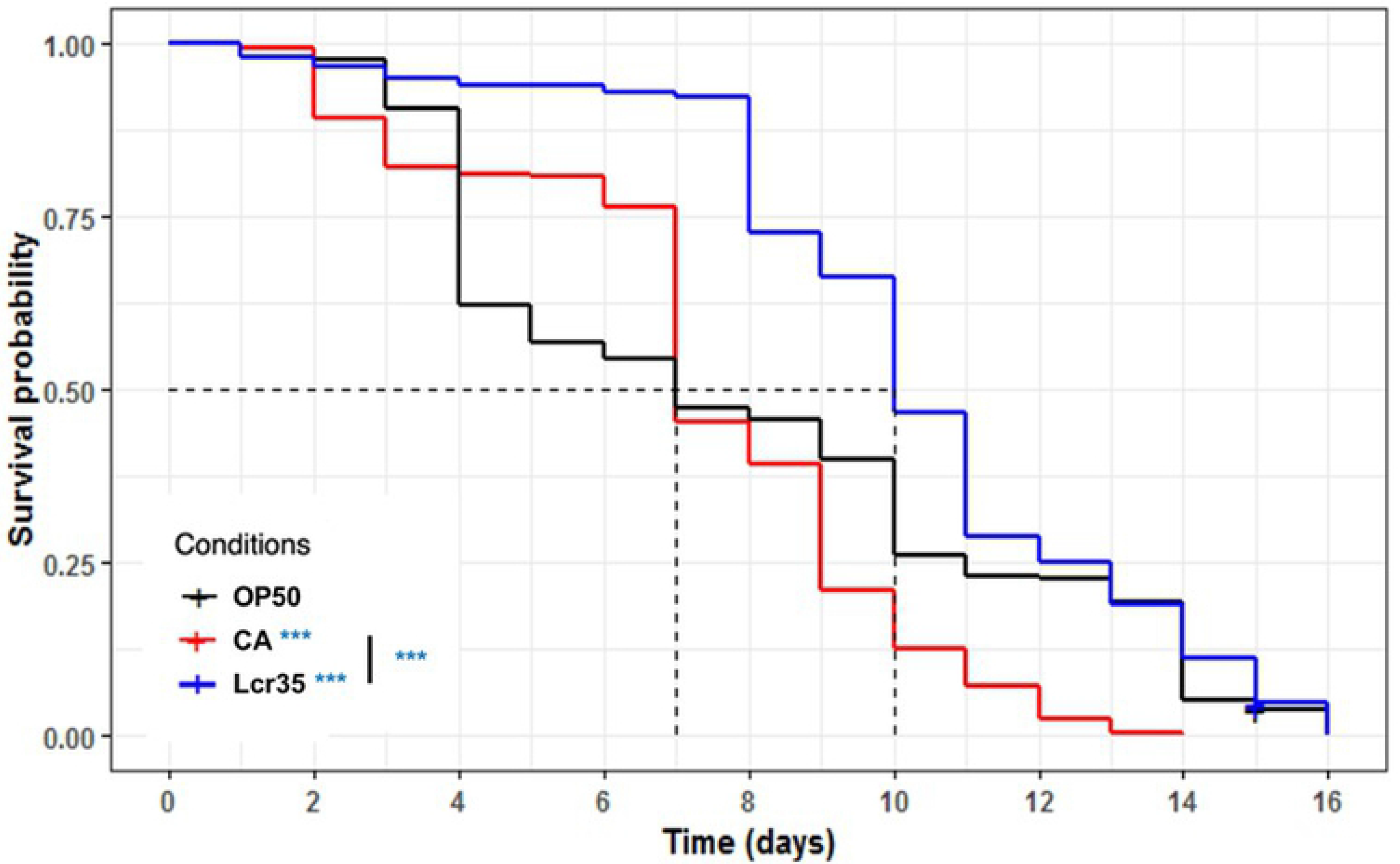
Influence of *Lactobacillus rhamnosus* Lcr35^®^ on lifespan of *C. elegans* wild-type N2 strain. Worms were fed with *E. coli* OP50 (n = 285) *C. albicans* ATCC 10231 (n = 242), and Lcr35^®^ (n = 278). Mean lifespan, where half of the population is dead, is represented on the abscissa. The asterisks indicate the *p-values* (log-rank test) with OP50 as a control (p < 0.05 : * ; p < 0.01 : ** ; p < 0.001 : ***).

#### 3.2.2 Lcr35^®^ does not modify *C. elegans* growth

The body size of Lcr35^®^ fed nematodes were compared to OP50-fed worms. Feeding worms with the probiotic strain did not significantly change in growth rate nor body size as they all reached their maximal length after three days (Fig 3).

**Fig 3:**
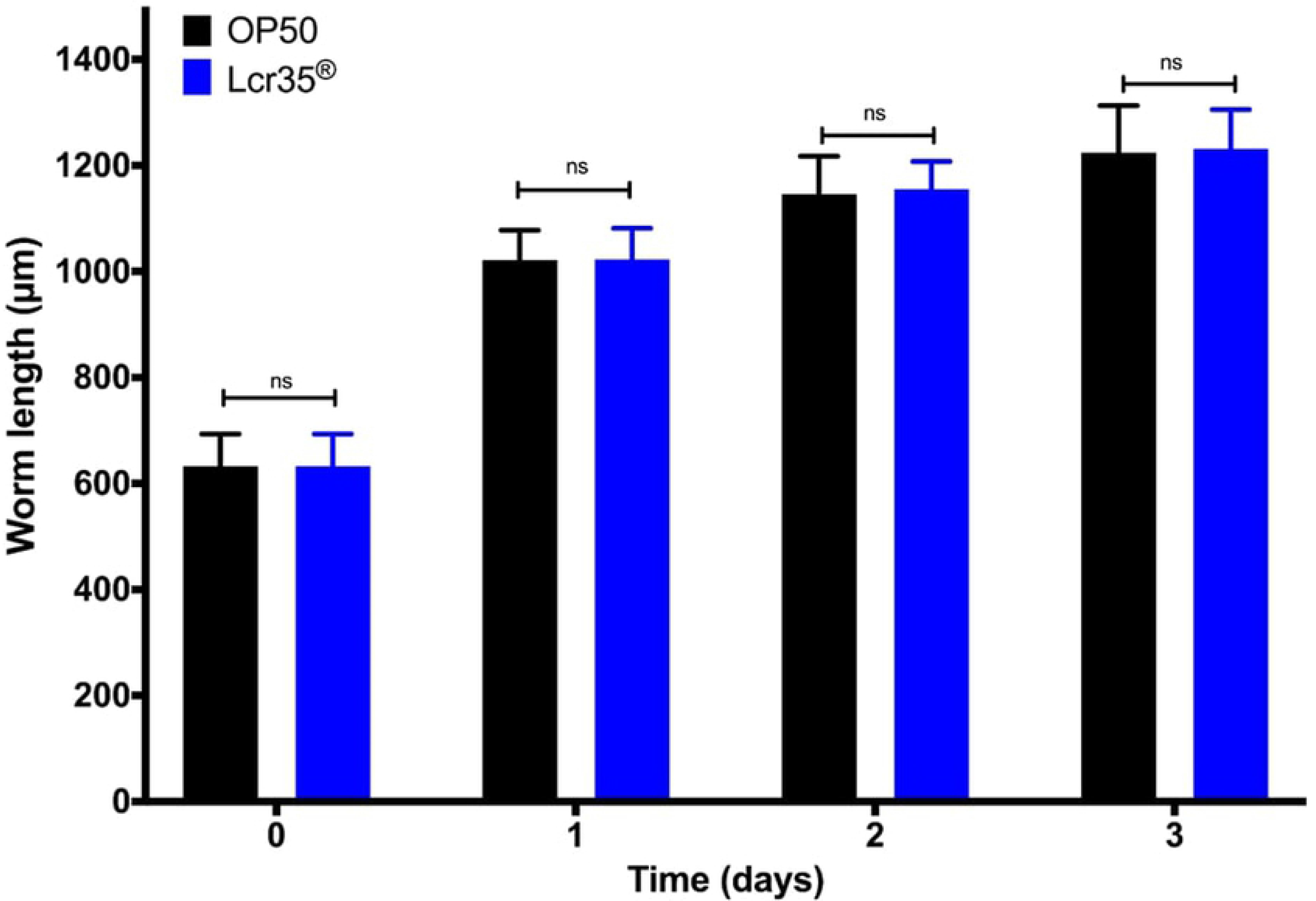
Growth of *C. elegans* (adult) on *E. coli* OP50 and on Lcr35^®^. All results are represented as means +/− standard deviations.

### 3.3 Effect of Lcr35^®^ preventive treatment on candidiasis

#### 3.3.1 Effect of Lcr35^®^ on *C. elegans* survival after *C. albicans* exposure

When *C. elegans* was sequentially exposed for 2 h to Lcr35^®^ prior being infected by *C. albicans*, the survival of the nematodes was increased significantly as the mean lifespan rised from 3 to 11 days (267% increase in survival) compared with that observed with *C. albicans* infection alone (p < 2.10^−16^). There was no significant difference between worms sequentially exposed to Lcr35^®^ and *C. albicans* and those exposed to Lcr35^®^ only (Fig 4) (p = 1). Similar results were obtained when the nematodes were exposed to the probiotic for 4 hours. In that case, we observed that Lcr35^®^ completely protected *C. elegans* from infection since there was no significant difference with the Lcr35^®^ control condition without infection (p = 0.4).

**Fig 4:**
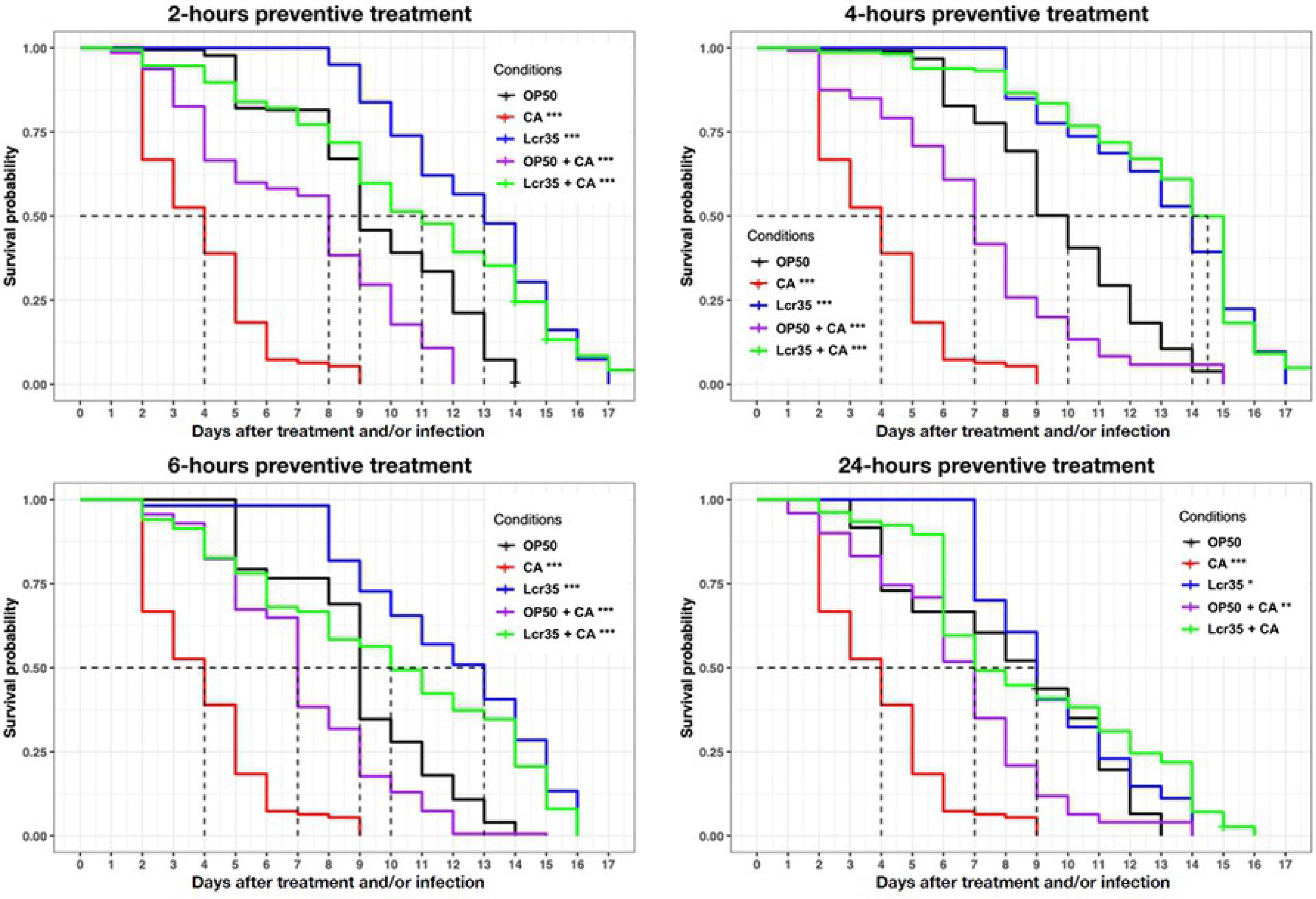
Preventive effects of Lcr35^®^ against *C. albicans* ATCC 10231. Mean survival, where half of the population is dead, is represented on the abscissa. The asterisks indicate the *p-values* (log-rank test) against OP50 (p < 0.05: * ; p < 0.01: ** ; p < 0.001 : ***). Infection duration: 2 hours; treatment duration: 2 hours (*E. coli* OP50 (n = 126); *C. albicans* ATCC 10231 (n = 424); Lcr35^®^ (n = 93); *C. albicans* + *E. coli* OP50 (n = 287); *C. albicans* + Lcr35^®^ (n = 224)) ; treatment duration: 4 hours (*E. coli* OP50 (n = 313); *C. albicans* ATCC 10231 (n = 424); Lcr35^®^ (n = 259); *C. albicans* + *E. coli* OP50 (n = 120); *C. albicans* + Lcr35^®^ (n = 164)); treatment duration: 6 hours (*E. coli* OP50 (n = 222); *C. albicans* ATCC 10231 (n = 424); Lcr35^®^ (n = 165); *C. albicans* + *E. coli* OP50 (n = 339); *C. albicans* + Lcr35^®^ (n = 300)); treatment duration: 24 hours (*E. coli* OP50 (n = 248); *C. albicans* ATCC 10231 (n = 424); Lcr35^®^ (n = 170); *C. albicans* + *E. coli* OP50 (n = 220); *C. albicans* + Lcr35^®^ (n = 183)).

For longer treatment times (6 and 24 hours), we observed a significant decrease of mean survival in the presence of Lcr35^®^ (condition 6 hours: p = 0.04, condition 24 hours: p <2.10^−16^) or Lcr35^®^ and *C. albicans* (condition 6 hours: p = 9.10^−13^, condition 24 hours: p < 2.10^−16^) compared to the treatment of 4 hours. Taken together, the results showed that the 4 hours probiotic treatment was the most protective against infection.

#### 3.3.2 Influence of Lcr35^®^ presence on *C. albicans* colonization of the worm’s gut

In order to determine whether the anti-*Candida* effects observed were due to the removal of the pathogen, colonization of the intestine of the nematode by *C. albicans* was observed by light microscopy. After three days of incubation in the presence of the pathogen, wild-type worms had an important colonization of the entire digestive tract (Fig 5A). However, it turned out that this strain of *C. albicans* was not able to form hyphae within the worm. We subsequently applied prophylactic treatment to the worms for 4 hours before infecting them with yeast. We observed that after a preventive treatment with the control OP50 (Fig 5B) or the probiotic Lcr35^®^ (Fig 5C), the yeast *C. albicans* was still detected in the digestive tract of the host.

**Fig 5:**
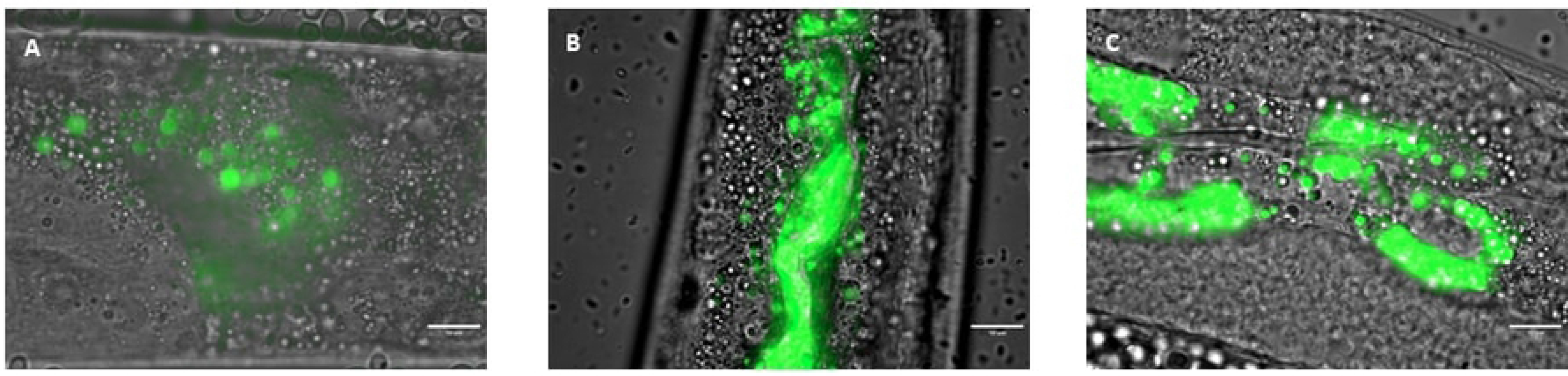
*C. albicans* colonization of *C. elegans*’s gut 72 hours (A) and after a 4-hour-prophylactic treatment with *E. coli* OP50 (B) or Lcr35^®^ (C) . The green color represents yeast labeled with rhodamine 123. Scale bar, 10 μm.

### 3.4 Mechanistic study

#### 3.4.1 Modulation of *C. elegans* genes expression induced by Lcr35^®^ and *C. albicans*

To elucidate the mechanisms involved in the action of Lcr35^®^ against *C. albicans*, we studied the expression of seven *C. elegans* genes (Table 3). We targeted three groups of genes: *daf-2* and *daf-16* (insulin signaling pathway) involved in host longevity and antipathogenicity, *sek-1* and *pmk-1* (p38 MAPK signaling pathway) which concern the immunity response as well as *abf-2*, *cnc-4* and *fipr-22* / *fipr-23* which encode for antimicrobial proteins. We noted that Lcr35^®^ tended to induce an overexpression of *daf-16* (p = 0.1635) while having no effect on *daf-2* (p = 0.2536) when *C. albicans* tended to induce an up-regulation of both genes (p = 0.1155 and p = 0.2396 respectively). We did not observe any expression modulation of *daf-2* nor *daf-16* using a preventive treatment with *E. coli* OP50 (p = 0.1258 and p = 0.1215) or with Lcr35^®^ (p = 0.1354 and p = 0.3021).

**Table 3:** Modulation of *C. elegans* GOIs expression induced by Lcr35^®^ and *C. albicans* in pure and sequential cultures in comparison with the control condition *E. coli* OP50 (alone). Genes were considered differentially expressed when the p-value was lower than 0.05 (*) or 0.01 (**) according to Fisher’s LSD test, and simultaneously when the expression change was of at least 2 times or 0.5 times.

| Conditions | Genes of interest |  |  |  |  |  |  |
| --- | --- | --- | --- | --- | --- | --- | --- |
|  | Insulin signaling pathway |  | p38 MAPK signaling pathway |  | Antimicrobials |  |  |
|  | *daf-2* | *daf-16* | *sek-1* | *pmk-1* | *abf-2* | *cnc-4* | *fipr-22* /<br>*fipr-23* |
| Lcr35<sup>®</sup> | 1.35 | 2.18 | 0.38 ** | 0.36 * | 1.70 | 3.39 | 0.61 |
| *C. albicans* | 2.48 | 3.31 | 3.21 * | 4.33 | 11.33 | 22.32 | 1.08 |
| *E. coli* OP50 + *C. albicans* | 1.82 | 0.53 | 0.37 * | 3.40 | 4.69 | 0.16 ** | 0.78 |
| Lcr35<sup>®</sup> + *C. albicans* | 0.69 | 1.74 | 0.31 ** | 1.15 | 1.61 | 0.41 * | 0.42 * |

The *sek-1* and *pmk-1* immunity genes were significantly downregulated in the presence of Lcr35^®^ by a 2.63-fold (p = 0.015) and 2.78-fold (p = 0.0149) while they were up-regulated by *C. albicans* 3.21-fold (p = 0.0247) and 4.33-fold (0.1618). Preventive treatment with *E. coli* OP50 repressed 2.70 times *sek-1* (0.37-fold with p = 0.0204) but tended to overexpress *pmk-1*. Preventive treatment with Lcr35^®^ had the same effect on *sek-1* (p = 0.0016) but induced no change on *pmk-1* expression (p = 0.8205). Finally, among the 3 antimicrobials encoding genes tested, only the expression of *cnc-4* seemed to be modulated in the presence of Lcr35^®^ with an overexpression (p = 0.1753). *C albicans* seemed also to induce overexpression of *abf-2* (p = 0.2213) and *cnc-4* (p = 0.3228) but interestingly, *fipr-22* / *fipr-23* (p = 0.8225) expression remained unchanged. Overexpression of *abf-2* (6.25-fold, p = 0.3158) and significant repression of *cnc-4* (p = 0.0088) were observed when *E. coli* OP50 was used as a preventive treatment. Using a Lc35^®^ preventive treatment, *cnc-4* and *fipr-22* / *fipr-23* were significantly repressed (p = 0.0396 and p = 0.0385 respectively).

#### 3.4.2 Influence of Lcr35^®^ and *C. albicans* on DAF-16 nuclear translocation

In order to further investigate the mechanisms involved in the anti-*C. albicans* effects of Lcr35^®^, we followed the nuclear translocation of DAF-16 / FOXO transcription factor using DAF-16::GFP strain. Whatever the incubation time, the worms did not show any translocation of DAF-16 while feeding with *E. coli* OP50 (Fig 6A). When Lcr35^®^ is used as food, we observed a nuclear translocation of the transcription factor, taking place gradually from 4 hours of incubation with a maximum of intensity in the nuclei after 6 hours. The distribution of DAF-16 was both cytoplasmic and nuclear (Fig 6B). When the nematode was fed exclusively with *C. albicans*, we observed a rapid nuclear translocation of the transcription factor after two hours of incubation in the presence of the pathogen (Fig 6C). This translocation was maintained throughout the experiment i.e. 76 hours.

**Fig 6:**
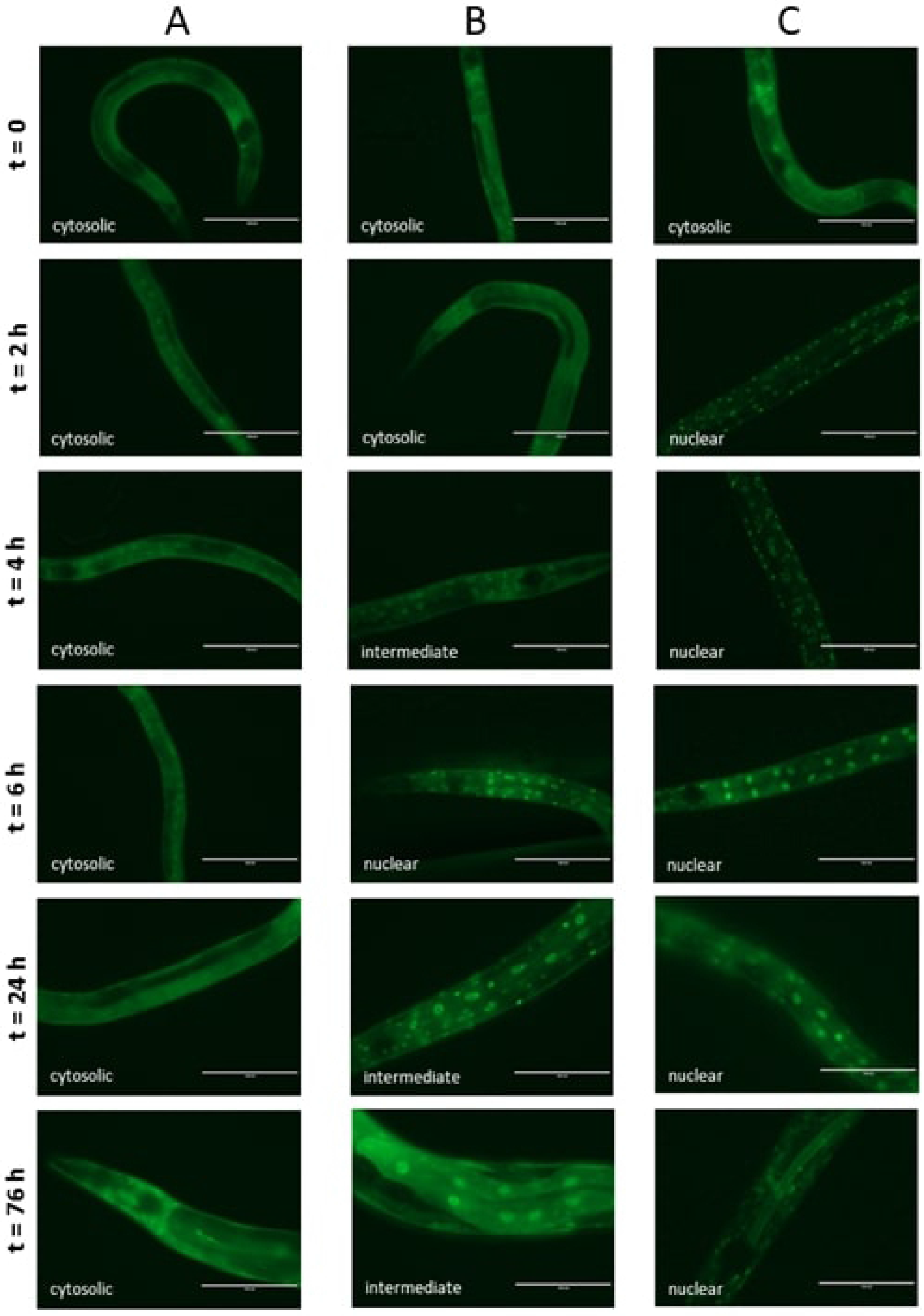
DAF-16 cellular localization in *C. elegans* transgenic strain TJ-356 (*daf-16p::daf-16a/b::GFP* +*rol-6(su1006)*) expressing DAF-16::GFP. Worms fed on OP50 (A), on Lcr35^®^ (B) and on *C. albicans* ATCC 10231 (C). Scale bar, 100 μm

#### 3.4.3 Effect of Lcr35^®^ preventive treatment on DAF-16 nuclear translocation

We investigated the effect of preventive treatment on the cellular localization of DAF-16 over time after infection by *C. albicans*. When nematodes were first fed with *E. coli* OP50 before being infected, DAF-16 was fully observed in the nuclei up to 4 hours after infection and then gradually translocated to be cytoplasmic after 24 hours (Fig 7A). Conversely, the worms first exposed to Lcr35^®^ and then to the pathogen showed a different response, the transcription factor was found only in the nuclei (Fig 7B).

**Fig 7:**
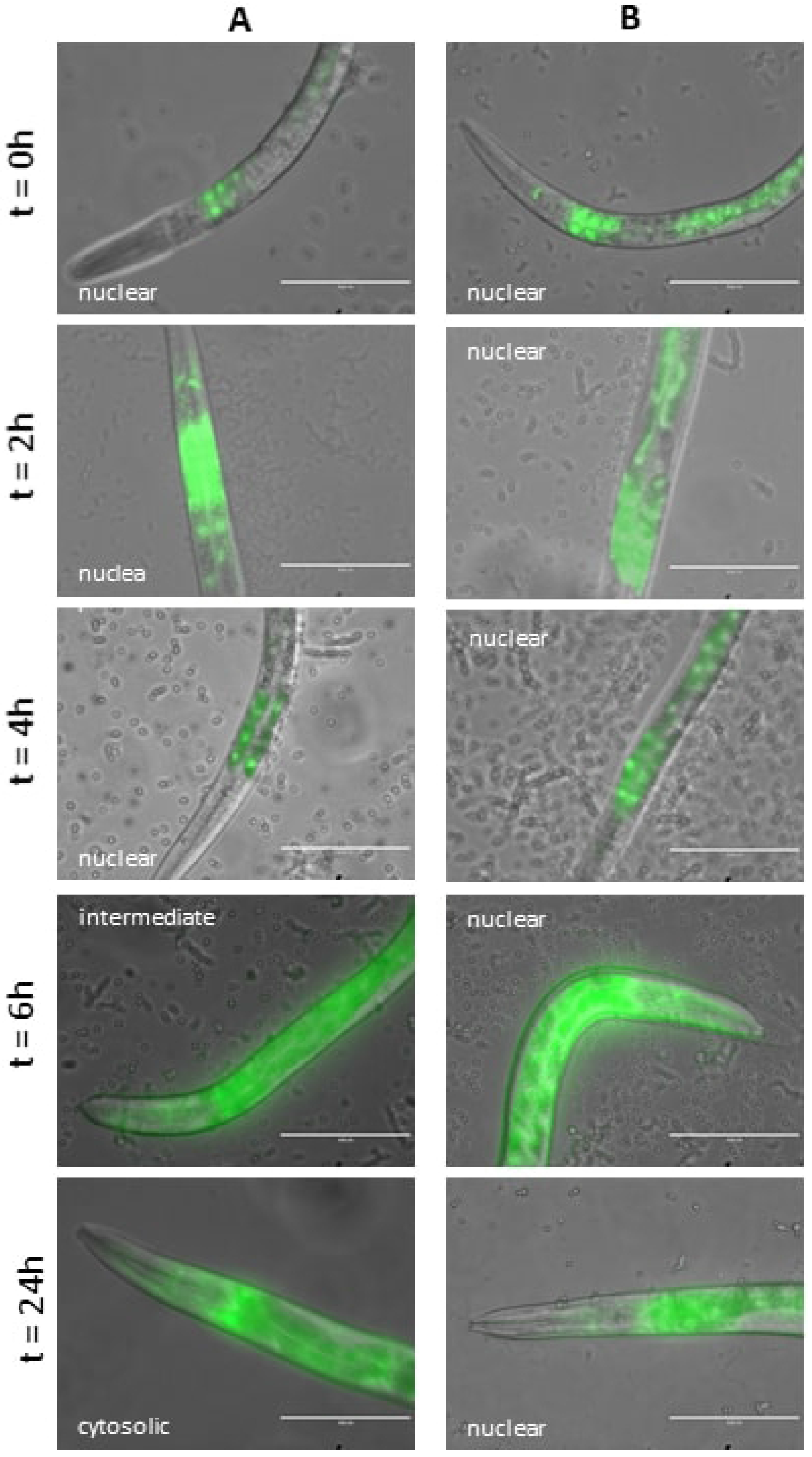
Effect of preventive approach on DAF-16 nuclear localization in *C. elegans* transgenic strain TJ-356 expressing DAF-16::GFP. Worms fed on OP50 + *C. albicans* (A) and on Lcr35^®^ + *C. albicans* (B). Scale bar, 100 μm

## 4 Discussion

Selection of microbial strains as probiotics is based on a combination of functional probiotic properties revealed first by classical basic *in vitro* testing. Beyond resistance to gastric pH or bile salts, the ability of the strain to adhere to epithelial cells is frequently studied since this represents a prerequisite for the mucosal colonization as part of the anti-pathogen activity. Adhesion is also a key parameter for pathogens since it allows them to release toxins and enzymes directly into the target cell, facilitating their dissemination (47). Nivoliez *et al.* showed that native probiotic strain Lcr35^®^ adhered rather weakly to the Caco-2 intestinal cells while the industrial formulation increases this capacity (24). We have further demonstrated here the ability of Lcr35^®^ to inhibit the growth of the pathogen *C. albicans* and the formation of a *Candida* biofilm on an intestinal cells monolayer *in vitro*. As described by Jankowska *et al.* (47), the low adherence of *L. rhamnosus* compared to *C. albicans* seems to reflect that competition for membrane receptors is not the only mechanism. It is probably related to the synthesis of antifungal effectors by the probiotic as well (47). Exopolysaccharides (EPS) secreted by certain lactobacilli have been shown to modify the surface properties (hydrophobicity) of microorganisms with direct consequences on their adhesion capacities (48). EPS have antifungal effect by inhibiting the growth of *C. albicans* but also its adhesion to epithelial cells. The surface polysaccharides of *L. rhamnosus* GG, one strain phylogenetically close to Lcr35, appear to interfere in the binding between the fungal lectin-like adhesins and host sugars or between the fungal cell wall carbohydrates and their epithelial adhesion receptor (49). A recent study has shown that purified fractions of exopolysaccharides also interfered with adhesion capacities of microorganisms (50). It would be interesting to assay the inhibitory properties of Lcr35^®^ EPS. But in order to fully understand the probiotic mechanisms, *in vitro* approaches are too limited. Moving to an *in vivo* approach is mandatory to better understand the interactions between microorganisms (probiotics and pathogens) and the host response.

*C. elegans* is considered as a powerful *in vivo* model for studying the pathogenicity of microorganisms (34,35,51–53) but also the antimicrobial properties of lactic acid bacteria (54,55). The nature of the nutrient source is an important parameter that has a great influence on the nematode’s physiology. Depending on the quality and quantity of food, the growth and body size, fertility and longevity of *C. elegans* are affected either positively or negatively (56,57). Regarding to worm growth, it appears that there is some disparity depending on the type of lactic acid bacteria used. It has been shown that *Bifidobacterium spp*. had no influence on the size of adult worms although their growth is slightly slowed down (58,59). *Lactobacillus spp.* by contrast usually result in lower growth rates but also lower sizes and are sometimes even lethal to the larvae (60,61). The mechanisms for explaining the longevity extension induced by lactic acid bacteria are not fully understood. Suggested by some authors, caloric restriction is known as a method of extending the lifespan of many taxa (62–64). In our case, similarly to the work of Komura *et al.*, it seems that it is not involved in the present case insofar as the growth of nematodes in the presence of the probiotic is strictly identical compared to *E. coli* OP50-fed worms (65).

After demonstrating the preventive effect of Lcr35 in the nematode, we decided to better understand the protective effect at the mechanistic level. In *C. elegans*, the insulin / IGF-1 signaling pathway is strongly involved in regulating the longevity and immunity of the animal. Signal transduction is mediated through DAF-16, a highly conserved FOXO transcription factor (66). Using the GFP fusion protein, we have shown that Lcr35^®^ induces translocation of DAF-16 to the nucleus, suggesting that DAF-16 is involved in the probiotic mechanisms of action of Lcr35^®^. According to several studies, the pro-longevity effect of probiotics linked to DAF-16 implements strain-dependent mechanisms involving different regulatory pathways such as the DAF-2 / DAF-16 insulin pathway (67) or the c-Jun N-terminal kinase JNK-1 / DAF-16 pathway (59). The absence of modulation of *daf-2* expression in the presence of Lcr35^®^ suggests that the DAF-2 / DAF-16 pathway is not involved and that it is rather the JNK signaling pathway. The involvement of these pathways needs to be followed at proteomic and phosphoproteomic levels in order to validate this hypothesis.

The yeast *C. albicans* is capable of inducing a severe infection in *C. elegans* causing a rapid death of the host and even after a very short contact time. This infection is first manifested by the colonization of the whole intestinal lumen by yeasts and then, in the case of a virulent strain, by the formation of hyphae piercing the cuticle of the nematode leading to its death (34,68). In addition, it has been shown that strains of *C. albicans* incapable of forming hyphae, such as SPT20 mutants, have a significantly reduced pathogenicity in *C. elegans* as well as in *Galleria mellonella* or *Mus musculus* models while still being lethal (37). In the nematode, it seems that the distention of the intestine caused by the accumulation of yeasts is one of the causes of the death of the animal (35). Recently, de Barros *et al.* (38) showed that *Lactobacillus paracasei* 28.4 had anti-*C. albicans* activity both *in vitro* and *in vivo* by inhibiting filamentation of yeast protecting the nematode. Although *C. albicans* ATCC 10231 is able to form hyphae during *in vitro* assay, it failed to kill *C. elegans* by filamentation (data not shown). Therefore, it is likely that Lcr35^®^ represses virulence factors in yeast other than filamentation.

From a mechanistic point of view, we can venture several hypotheses that can explain the anti-*C. albicans* properties of Lcr35^®^ in the nematode: a direct interaction between the two microorganisms as well as an immunomodulation of the host by the probiotic. In the first case, it was demonstrated the inhibitory capacity of Lcr35^®^ with respect to the pathogen during co-culture (24) and on mammalian cells monolayers (this study). This inhibition may be due to nutrient competition (*i.e.* glycogen consumption) or to the production of toxic metabolites against the yeast (24). We have shown that even after a preventive treatment with the probiotic, the digestive tract of the nematode is colonized by the pathogen without showing a pathological state. This suggests that Lcr35^®^ induced repression of virulence factors in *C. albicans* as this has been shown by De Barros *et al*. (38). In the second case, an *in vitro* study on human dendritic cells revealed that Lcr35^®^ induced a large dose-dependent modulation in the expression of genes mainly involved in the immune response but also in the expression of CD, HLA and TLR membrane proteins. Highly conserved and found in *C. elegans*, TLR also play a role in the antipathogenic response of the nematode by activating the p38 MAPK pathway. Kim and Mylonakis (2012) showed that *tir-1* was involved in the probiotic mechanism of *L. acidophilus* NCFM (55). A pro-inflammatory effect has also been shown through cytokine secretion such as IL-1β, IL-12, TNFα. However, this immunomodulation takes place only in the presence of high concentration of Lcr35^®^ (69). In *C. elegans*, DAF-16 is closely related to mammalian FOXO3a, a transcription factor involved the inflammatory process (70). Therefore, activation of DAF-16 by Lcr35^®^ can be interpreted as the establishment of an inflammatory response in the host and allowing it to survive an infection. In our study, we observed that the duration of the Lcr35^®^ treatment influences the preventive anti-*Candida* effect on nematode lifespan suggesting that the quantity of Lcr35^®^ ingested and/or treatment period of time may have an impact on the efficiency of the treatment. A thorough transcriptional study is interesting to characterize the deleterious effect of an increase in the dose of probiotics administered. We demonstrate that Lcr35^®^ induces a transcriptional response in the host by activating the transcription factor DAF-16 and repressing the p38 MAPK signaling pathway, including in the presence of *C. albicans*. We also observe the repression of the genes encoding for antimicrobials when the fungal infection was preceded by the probiotic treatment. The work of Pukkila-Worley *et al.* (35) demonstrated that *C. albicans* induced a fast antifungal response in the host inducing the secretion of antimicrobials such as *abf-2*, *cnc-4*, *cnc-7*, *fipr-22* and *fipr-23*. With the exception of *abf-2*, all these genes are under the control of PMK-1 whose inactivation makes the nematode susceptible to infection. In our study, we showed an Lcr35^®^ preventive treatment induced a down regulation of *cnc-4*, *fipr-22* and *fipr-23* genes while *pmk-1* remained unchanged compared to the control condition. The absence of overexpression of these genes in the presence of *C. albicans* after a pre-exposure with Lcr35^®^ suggests again that the probiotic inhibits the yeast virulence obviating the establishment of a defense mechanism by the host. Similar results have also been observed with *Salmonella* Enteritidis where the authors hypothesize that the probiotics used induce immunotolerance in the nematode rather than the synthesis of antimicrobials (58). The use of *C. elegans* mutants or RNAi could be further considered to decipher the signaling and regulation mechanisms.

## 5 Conclusion

This study demonstrates the preventive anti-*C. albicans* properties of Lcr35^®^ using both *in vitro* and *in vivo* preclinical models. The probiotic strain inhibits the growth of the pathogenic yeast and its ability to form biofilm on intestinal cells *in vitro*. Lcr35^®^ allows a protection of the host *C. elegans* against infection despite the presence of *C. albicans* in its gut. Lcr35^®^ during *C. albicans* infection seems to induce a decrease in the immune response of the nematode (downregulation of *sek-1*, *pmk-1*, *abf-2*, *cnc-4* and *fipr-22* / *23*). Extra studies on *C. elegans* whole transcriptome modulation by Lcr35^®^ would be interesting to further reveal other mechanisms involved. The study of the yeast virulence genes modulation induced by Lcr35^®^ could be very informative about complex mechanisms of the probiotic mechanisms of action. Also, in a second phase, the realization of a comparative study between Lcr35^®^ and other *Lactobacillus* strains (*L. rhamnosus*, *L. casei*, *L. paracasei*) could be of interest to determine the degree of strain-dependence of our results.

## 6 Acknowledgments

Some strains were provided by the CGC, which is funded by NIH Office of Research Infrastructure Programs (P40 OD010440).

We thank Jonathan Heuzé, Muriel Théret and Jeanne Riom, trainees at the UMRF 0545 (UCA/INRA) for their involvement in the implementation of several experiments. We thank greatly all those who took part in the writing of this article.

## 8 Author Contributions

CP, MG and OC conceived and planned the experiments. CP, TS, PV, MG and OC carried out the experiments with help from MB and SB. CP wrote the manuscript. PV, MB, CD, CC and SB provided critical feedback. AN and SB supervised the project.

## References

1. Cauchie M, Desmet S, Lagrou K. *Candida* and its dual lifestyle as a commensal and a pathogen. Res Microbiol [Internet]. 2017 Nov [cited 2018 Sep 5];168(9–10):802–10. Available from: https://linkinghub.elsevier.com/retrieve/pii/S0923250817300402

2. Neville BA, D’enfert C, Bougnoux M-E. *Candida albicans* commensalism in the gastrointestinal tract. FEMS Yeast Res [Internet]. 2015 [cited 2018 Sep 5];15:81. Available from: https://unite.ut.ee/

3. Mayer FL, Wilson D, Hube B. *Candida albicans* pathogenicity mechanisms. Vol. 4, Virulence. 2013. p. 119–28.

4. Kadosh D, Antonio S. Control of *Candida albicans* morphology and pathogenicity by post-transcriptional mechanisms. Cell Mol Life Sci. 2017;73(22):4265–78.

5. Wächtler B, Wilson D, Haedicke K, Dalle F, Hube B. From attachment to damage: Defined genes of *Candida albicans* mediate adhesion, invasion and damage during interaction with oral epithelial cells. Munro C, editor. PLoS One [Internet]. 2011 Feb 23 [cited 2018 Sep 10];6(2):e17046. Available from: http://www.ncbi.nlm.nih.gov/pubmed/21407800

6. Farmakiotis D, Kontoyiannis DP. Epidemiology of antifungal resistance in human pathogenic yeasts: current viewpoint and practical recommendations for management. Int J Antimicrob Agents [Internet]. 2017 Sep [cited 2018 Sep 5];50(3):318–24. Available from: https://linkinghub.elsevier.com/retrieve/pii/S0924857917302364

7. Sanguinetti M, Posteraro B, Lass-Flörl C. Antifungal drug resistance among *Candida* species: Mechanisms and clinical impact. Mycoses [Internet]. 2015 Jun [cited 2018 Sep 5];58(S2):2–13. Available from: http://www.ncbi.nlm.nih.gov/pubmed/26033251

8. Scorzoni L, de Paula E Silva ACA, Marcos CM, Assato PA, de Melo WCMA, de Oliveira HC, et al. Antifungal Therapy: New Advances in the Understanding and Treatment of Mycosis. Front Microbiol [Internet]. 2017 [cited 2018 Sep 5];8:36. Available from: http://www.ncbi.nlm.nih.gov/pubmed/28167935

9. Wheeler ML, Limon JJ, Bar AS, Leal CA, Gargus M, Tang J, et al. Immunological Consequences of Intestinal Fungal Dysbiosis. Cell Host Microbe [Internet]. 2016;19(6):865–73. Available from: http://dx.doi.org/10.1016/j.chom.2016.05.003

10. Hu H-J, Zhang G-Q, Zhang Q, Shakya S, Li Z-Y. Probiotics Prevent *Candida* Colonization and Invasive Fungal Sepsis in Preterm Neonates: A Systematic Review and Meta-Analysis of Randomized Controlled Trials. Pediatr Neonatol [Internet]. 2017 Apr [cited 2018 Sep 5];58(2):103–10. Available from: http://linkinghub.elsevier.com/retrieve/pii/S1875957216301401

11. Matsubara VH, Bandara HMHN, Mayer MPA, Samaranayake LP. Probiotics as Antifungals in Mucosal Candidiasis. 2016 [cited 2018 Sep 5]; Available from: https://academic.oup.com/cid/article-abstract/62/9/1143/1745140

12. Agrawal S, Rao S, Patole S. Probiotic supplementation for preventing invasive fungal infections in preterm neonates - a systematic review and meta-analysis. Mycoses [Internet]. 2015 Nov 1 [cited 2018 Sep 5];58(11):642–51. Available from: http://doi.wiley.com/10.1111/myc.12368

13. FAO, WHO. Health and Nutritional Properties of Probiotics in Food including Powder Milk with Live Lactic Acid Bacteria. Food Nutr Pap [Internet]. 2001 [cited 2016 Jun 14]; Available from: http://www.crcnetbase.com/doi/abs/10.1201/9781420009613.ch16

14. Fijan S. Microorganisms with Claimed Probiotic Properties: An Overview of Recent Literature. Int J Environ Res Public Heal Int J Environ Res Public Heal Int J Environ Res Public Heal [Internet]. 2014 [cited 2017 May 13];11:4745–67. Available from: www.mdpi.com/journal/ijerph

15. Olle B. Medicines from microbiota. Nat Biotechnol [Internet]. 2013 Apr 5 [cited 2017 Mar 31];31(4):309–15. Available from: http://www.nature.com/doifinder/10.1038/nbt.2548

16. Coudeyras S, Jugie G, Vermerie M, Forestier C. Adhesion of human probiotic *Lactobacillus rhamnosus* to cervical and vaginal cells and interaction with vaginosis-associated pathogens. Infect Dis Obstet Gynecol [Internet]. 2008 Jan 27 [cited 2018 Sep 10];2008:549640. Available from: http://www.ncbi.nlm.nih.gov/pubmed/19190778

17. Coudeyras S, Marchandin H, Fajon C, Forestier C. Taxonomic and strain-specific identification of the probiotic strain *Lactobacillus rhamnosus* 35 within the *Lactobacillus casei* group. Appl Environ Microbiol [Internet]. 2008 May [cited 2018 Sep 10];74(9):2679–89. Available from: http://www.ncbi.nlm.nih.gov/pubmed/18326671

18. Forestier C, De Champs C, Vatoux C, Joly B. Probiotic activities of *Lactobacillus casei rhamnosus*: *in vitro* adherence to intestinal cells and antimicrobial properties. Res Microbiol [Internet]. 2001 Mar [cited 2018 Sep 10];152(2):167–73. Available from: http://www.ncbi.nlm.nih.gov/pubmed/11316370

19. de Champs C, Maroncle N, Balestrino D, Rich C, Forestier C. Persistence of colonization of intestinal mucosa by a probiotic strain, *Lactobacillus casei* subsp. *rhamnosus* Lcr35, after oral consumption. J Clin Microbiol [Internet]. 2003 Mar [cited 2018 Sep 10];41(3):1270–3. Available from: http://www.ncbi.nlm.nih.gov/pubmed/12624065

20. Petricevic L, Witt A. The role of *Lactobacillus casei rhamnosus* Lcr35 in restoring the normal vaginal flora after antibiotic treatment of bacterial vaginosis. BJOG An Int J Obstet Gynaecol [Internet]. 2008 Oct [cited 2016 Jun 14];115(11):1369–74. Available from: http://doi.wiley.com/10.1111/j.1471-0528.2008.01882.x

21. Muller C, Mazel V, Dausset C, Busignies V, Bornes S, Nivoliez A, et al. Study of the *Lactobacillus rhamnosus* Lcr35^®^ properties after compression and proposition of a model to predict tablet stability. Eur J Pharm Biopharm. 2014;88(3):787–94.

22. Nivoliez A, Veisseire P, Alaterre E, Dausset C, Baptiste F, Camarès O, et al. Influence of manufacturing processes on cell surface properties of probiotic strain *Lactobacillus rhamnosus* Lcr35^®^. Appl Microbiol Biotechnol [Internet]. 2015 [cited 2017 Jan 1];99(1):399–411. Available from: http://link.springer.com/10.1007/s00253-014-6110-z

23. Dausset C, Patrier S, Gajer P, Thoral C, Lenglet Y, Cardot JM, et al. Comparative phase I randomized open-label pilot clinical trial of Gynophilus^®^ (Lcr regenerans®) immediate release capsules versus slow release muco-adhesive tablets. Eur J Clin Microbiol Infect Dis [Internet]. 2018 [cited 2019 Apr 2];37(10):1869–80. Available from: https://doi.org/10.1007/s10096-018-3321-8

24. Nivoliez A, Camares O, Paquet-Gachinat M, Bornes S, Forestier C, Veisseire P. Influence of manufacturing processes on *in vitro* properties of the probiotic strain *Lactobacillus rhamnosus* Lcr35^®^. J Biotechnol. 2012;160(3–4):236–41.

25. Isolauri E, Kirjavainen P V, Salminen S. Probiotics: a role in the treatment of intestinal infection and inflammation? Gut [Internet]. 2002;50(Supplement 3):iii54–iii59. Available from: http://gut.bmj.com/cgi/doi/10.1136/gut.50.suppl_3.iii54

26. do Carmo MS, Santos C itapary dos, Araújo MC, Girón JA, Fernandes ES, Monteiro-Neto V. Probiotics, mechanisms of action, and clinical perspectives for diarrhea management in children. Food Funct [Internet]. 2018;9(10):5074–95. Available from: http://dx.doi.org/10.1039/c8fo00376a

27. Coudeyras S, Forestier C. Microbiote et probiotiques : impact en santé humaine. Can J Microbiol [Internet]. 2010 [cited 2018 Jan 30];56(8):611–50. Available from: http://www.nrcresearchpress.com/doi/pdfplus/10.1139/W10-052

28. Lacroix C, de Wouters T, Chassard C. Integrated multi-scale strategies to investigate nutritional compounds and their effect on the gut microbiota. Curr Opin Biotechnol [Internet]. 2015 [cited 2017 Apr 30];32:149–55. Available from: http://dx.doi.org/10.1016/j.copbio.2014.12.009

29. Vinderola G, Gueimonde M, Gomez-Gallego C, Delfederico L, Salminen S. Correlation between *in vitro* and *in vivo* assays in selection of probiotics from traditional species of bacteria. Trends Food Sci Technol [Internet]. 2017;68:83–90. Available from: http://dx.doi.org/10.1016/j.tifs.2017.08.005

30. Montoro BP, Benomar N, Lerma LL, Gutiérrez SC, Gálvez A, Abriouel H. Fermented aloreña table olives as a source of potential probiotic *Lactobacillus pentosus* strains. Front Microbiol. 2016;7(OCT).

31. Roselli M, Finamore A, Britti MS, Mengheri E. Probiotic bacteria *Bifidobacterium animalis* MB5 and *Lactobacillus rhamnosus* GG protect intestinal Caco-2 cells from the inflammation-associated response induced by enterotoxigenic *Escherichia coli* K88. Br J Nutr [Internet]. 2006;95(06):1177. Available from: http://www.journals.cambridge.org/abstract_S0007114506001589

32. Papadimitriou K, Zoumpopoulou G, Foligné B, Alexandraki V, Kazou M, Pot B, et al. Discovering probiotic microorganisms: *In vitro*, *in vivo*, genetic and omics approaches. Front Microbiol. 2015;6(FEB):1–28.

33. Lai CH, Chou CY, Ch’ang LY, Liu CS, Lin W. Identification of novel human genes evolutionarily conserved in *Caenorhabditis elegans* by comparative proteomics. Genome Res [Internet]. 2000 May [cited 2018 Sep 10];10(5):703–13. Available from: http://www.ncbi.nlm.nih.gov/pubmed/10810093

34. Pukkila-Worley R, Peleg AY, Tampakakis E, Mylonakis E. *Candida albicans* hyphal formation and virulence assessed using a *Caenorhabditis elegans* infection model. Eukaryot Cell [Internet]. 2009 [cited 2018 Feb 7];8(11):1750–8. Available from: http://ec.asm.org/content/8/11/1750.full.pdf

35. Pukkila-Worley R, Ausubel FM, Mylonakis E. *Candida albicans* infection of *Caenorhabditis elegans* induces antifungal immune defenses. PLoS Pathog. 2011;7(6).

36. Alves V de S, Mylonakis E. The eIF2 kinase Gcn2 modulates Candida albicans virulence to Caenorhabditis elegans. Clin Microbiol Infect Dis [Internet]. 2018;3(2):1–4. Available from: https://www.oatext.com/the-eif2-kinase-gcn2-modulates-candida-albicans-virulence-to-caenorhabditis-elegans.php

37. Tan X, Fuchs BB, Wang Y, Chen W, Yuen GJ, Chen RB, et al. The role of *Candida albicans* SPT20 in filamentation, biofilm formation and pathogenesis. PLoS One. 2014;9(4):1–10.

38. de Barros PP, Scorzoni L, Ribeiro F de C, Fugisaki LR de O, Fuchs BB, Mylonakis E, et al. *Lactobacillus paracasei* 28.4 reduces *in vitro* hyphae formation of *Candida albicans* and prevents the filamentation in an experimental model of *Caenorhabditis elegans*. Microb Pathog [Internet]. 2018;117(November 2017):80–7. Available from: https://doi.org/10.1016/j.micpath.2018.02.019

39. Pinto M, Robineleon S, Appay MD, Kedinger M, Triadou N, Dussaulx E, et al. Enterocyte-like differentiation and polarization of the human colon carcinoma cell line Caco-2 in culture. Biol Cell [Internet]. 1983 Jan 1 [cited 2018 Oct 10];47:323–30. Available from: https://www.scienceopen.com/document?vid=07f3fdcd-c23c-47d4-ad63-105346ef5453

40. Brenner S. The genetics of *Caenorhabditis elegans*. Genetics. 1974;77(1):71–94.

41. Mörck C, Pilon M. *C. elegans* feeding defective mutants have shorter body lengths and increased autophagy. BMC Dev Biol [Internet]. 2006 [cited 2018 Jan 30];6. Available from: https://www.ncbi.nlm.nih.gov/pmc/articles/PMC1559592/pdf/1471-213X-6-39.pdf

42. Hellemans J, Mortier G, De Paepe A, Speleman F, Vandesompele J. qBase relative quantification framework and software for management and automated analysis of real-time quantitative PCR data. 2007 [cited 2017 Jun 14];8(2). Available from: http://download.springer.com/static/pdf/804/art%253A10.1186%252Fgb-2007-8-2-r19.pdf?originUrl= http%3A%2F%2Fgenomebiology.biomedcentral.com%2Farticle%2F10.1186%2Fgb-2007-8-2-r19&token2=exp=1497424427~acl=%2Fstatic%2Fpdf%2F804%2Fart%25253A10.1186%25252Fgb-2007-8-2-r19.pdf*~hmac=5b660aaa729e6c840581e3f1a59b3a251732c03c6248935dd629971b9f7036f6

43. R Core Team. R: A language and Environment for Statistical Computing [Internet]. Vienna, Austria: R Foundation for Statistical Computing; 2018. Available from: https://www.r-project.org/

44. Therneau TM. _A Package for Survival Analysis in S_. 2015.

45. Kassambara A, Kosinski M. survminer: Drawing Survival Curves using “ggplot2.” 2017.

46. Fatima S, Haque R, Jadiya P, Shamsuzzama, Kumar L, Nazir A. Ida-1, the *Caenorhabditis elegans* orthologue of mammalian diabetes autoantigen IA-2, potentially acts as a common modulator between Parkinson’s disease and diabetes: Role of Daf-2/Daf-16 insulin like signalling pathway. PLoS One. 2014;9(12).

47. Jankowska A, Laubitz D, Antushevich H, Zabielski R, Grzesiuk E. Competition of *Lactobacillus paracasei* with *Salmonella enterica* for adhesion to Caco-2 cells. J Biomed Biotechnol. 2008;2008(1).

48. Nowak A, Motyl I, Śliżewska K, Libudzisz Z, Klewicka E. Adherence of probiotic bacteria to human colon epithelial cells and inhibitory effect against enteric pathogens – *In vitro* study. Int J Dairy Technol. 2016;69(4):532–9.

49. Allonsius CN, van den Broek MFL, De Boeck I, Kiekens S, Oerlemans EFM, Kiekens F, et al. Interplay between *Lactobacillus rhamnosus* GG and *Candida* and the involvement of exopolysaccharides. Microb Biotechnol. 2017;10(6):1753–63.

50. Ruas-Madiedo P, Gueimonde M, Margolles A, de los Reyes-Gavilan CG, Salminen S. Exopolysaccharides Produced by Probiotic Strains Modify the Adhesion of Probiotics and Enteropathogens to Human Intestinal Mucus. J Food Prot [Internet]. 2006;69(8):2011–5. Available from: http://jfoodprotection.org/doi/abs/10.4315/0362-028X-69.8.2011

51. Irazoqui JE, Troemel ER, Feinbaum RL, Luhachack LG, Cezairliyan BO, Ausubel FM. Distinct pathogenesis and host responses during infection of *C. elegans* by *P. aeruginosa* and *S. aureus*. PLoS Pathog. 2010;6(7):1–24.

52. Wu K, Conly J, McClure JA, Elsayed S, Louie T, Zhang K. *Caenorhabditis elegans* as a host model for community-associated methicillin-resistant *Staphylococcus aureus*. Clin Microbiol Infect. 2010;16(3):245–54.

53. Souza ACR, Fuchs BB, Alves V de S, Jayamani E, Colombo AL, Mylonakis E. Pathogenesis of the *Candida parapsilosis* complex in the model host *Caenorhabditis elegans*. Genes (Basel). 2018;9(8).

54. Oh S, Park MR, Ryu S, Maburutse BE, Oh NS, Kim SH, et al. Probiotic *Lactobacillus fermentum* strain JDFM216 stimulates the longevity and immune response of *Caenorhabditis elegans* through a nuclear hormone receptor. Sci Rep [Internet]. 2018 [cited 2019 Jan 3];8(1):7441. Available from: www.nature.com/scientificreports/

55. Kim Y, Mylonakis E. *Caenorhabditis elegans* immune conditioning with the probiotic bacterium *Lactobacillus acidophilus* strain ncfm enhances gram-positive immune responses. Infect Immun. 2012;80(7):2500–8.

56. Yu L, Yan X, Ye C, Zhao H, Chen X, Hu F, et al. Bacterial respiration and growth rates affect the feeding preferences, brood size and lifespan of *Caenorhabditis elegans*. PLoS One. 2015;10(7):1–13.

57. So S, Miyahara K, Ohshima Y. Control of body size in *C. elegans* dependent on food and insulin/IGF-1 signal. Genes to Cells [Internet]. 2011 [cited 2018 Sep 23];16(6):639–51. Available from: https://onlinelibrary.wiley.com/doi/pdf/10.1111/j.1365-2443.2011.01514.x

58. Ikeda T, Yasui C, Hoshino K, Arikawa K, Nishikawa Y. Influence of lactic acid bacteria on longevity of *Caenorhabditis elegans* and host defense against *Salmonella enterica* serovar Enteritidis. Appl Environ Microbiol. 2007;73(20):6404–9.

59. Zhao L, Zhao Y, Liu R, Zheng X, Zhang M, Guo H, et al. The transcription factor DAF-16 is essential for increased longevity in *C. elegans* Exposed to *Bifidobacterium longum* BB68. Sci Rep [Internet]. 2017;7(1):7408. Available from: http://www.nature.com/articles/s41598-017-07974-3

60. Zanni E, Laudenzi C, Schifano E, Palleschi C, Perozzi G, Uccelletti D, et al. Impact of a complex food microbiota on energy metabolism in the model organism *Caenorhabditis elegans*. Biomed Res Int. 2015;2015.

61. Guantario B, Zinno P, Schifano E, Roselli M, Perozzi G, Palleschi C, et al. *In Vitro* and *in Vivo* selection of potentially probiotic lactobacilli from nocellara del belice table olives. Front Microbiol. 2018;9(MAR):595.

62. Phelan JP, Rose MR. Why dietary restriction substantially increases longevity in animal models but won’t in humans. Ageing Res Rev. 2005;4(3):339–50.

63. Smith ED, Kaeberlein TL, Lydum BT, Sager J, Welton KL, Kennedy BK, et al. Age- and calorie-independent life span extension from dietary restriction by bacterial deprivation in *Caenorhabditis elegans*. BMC Dev Biol. 2008;8:1–13.

64. Heestand BN, Shen Y, Liu W, Magner DB, Storm N, Meharg C, et al. Dietary Restriction Induced Longevity Is Mediated by Nuclear Receptor NHR-62 in *Caenorhabditis elegans*. PLoS Genet. 2013;9(7).

65. Komura T, Ikeda T, Yasui C, Saeki S, Nishikawa Y. Mechanism underlying prolongevity induced by bifidobacteria in *Caenorhabditis elegans*. Biogerontology. 2013;14(1):73–87.

66. Tullet JMA. DAF-16 target identification in *C. elegans*: past, present and future. Biogerontology [Internet]. 2015;16(2):221–34. Available from: http://link.springer.com/10.1007/s10522-014-9527-y

67. Grompone G, Martorell P, Llopis S, González N, Genovés S, Mulet AP, et al. Anti-Inflammatory *Lactobacillus rhamnosus* CNCM I-3690 Strain Protects against Oxidative Stress and Increases Lifespan in *Caenorhabditis elegans*. PLoS One. 2012;7(12).

68. Breger J, Fuchs BB, Aperis G, Moy TI, Ausubel FM, Mylonakis E. Antifungal chemical compounds identified using a *C. elegans* pathogenicity assay. PLoS Pathog. 2007;3(2):0168–78.

69. Evrard B, Coudeyras S, Dosgilbert A, Charbonnel N, Alamé J, Tridon A, et al. Dose-dependent immunomodulation of human dendritic cells by the probiotic *Lactobacillus rhamnosus* Lcr35. PLoS One. 2011;6(4):1–12.

70. Singh V, Aballay A. Regulation of DAF-16-mediated Innate Immunity in *Caenorhabditis elegans*. J Biol Chem [Internet]. 2009 Dec 18 [cited 2018 Dec 14];284(51):35580–7. Available from: http://www.ncbi.nlm.nih.gov/pubmed/19858203

71. Semple JI, Garcia-Verdugo R, Lehner B. Rapid selection of transgenic *C. elegans* using antibiotic resistance. Nat Methods [Internet]. 2010 Sep 22 [cited 2017 Apr 13];7(9):725–7. Available from: http://www.ncbi.nlm.nih.gov/pubmed/20729840

72. Hoogewijs D, Houthoofd K, Matthijssens F, Vandesompele J, Vanfleteren JR. Selection and validation of a set of reliable reference genes for quantitative *sod* gene expression analysis in *C. elegans*. BMC Mol Biol [Internet]. 2008 Jan 22 [cited 2017 Apr 13];9:9. Available from: http://www.ncbi.nlm.nih.gov/pubmed/18211699

73. Nakagawa H, Shiozaki T, Kobatake E, Hosoya T, Moriya T, Sakai F, et al. Effects and mechanisms of prolongevity induced by *Lactobacillus gasseri* SBT2055 in *Caenorhabditis elegans*. Aging Cell. 2016;15(2):227–36.

